# A Modular T7 RNAP Expression Architecture for Orthogonal Multigene Expression in Yeast

**DOI:** 10.64898/2026.06.19.733325

**Authors:** Eunha Jeon, Sujin Hong, Seung-Gyun Woo, Dae-Hee Lee

**Author notes:** These authors contributed equally to this work (Eunha Jeon, Sujin Hong). For correspondence: Dae-Hee Lee.

## Abstract

Orthogonal gene expression systems based on bacteriophage-derived T7 RNA polymerase (T7 RNAP) offer a promising strategy for decoupling engineered transcription from host regulatory networks. Although recent advances have enabled productive T7 RNAP-mediated expression in *Saccharomyces cerevisiae*, strategies for independently controlling multiple genes within a shared T7 transcriptional framework remain limited. Here, we developed a modular pYTK-compatible T7 RNAP expression architecture that separates overall transcriptional capacity from gene-specific control of protein output. We established interchangeable T7 promoter, terminator, polyadenylation signal, and 5’ untranslated region (UTR) parts for combinatorial assembly. Analysis of expression cassette dosage showed that transcript abundance and expression output scale with cassette copy number, with high-copy constructs producing transcript levels comparable to those driven by the *GAL1* promoter. To enable gene-specific tuning, we constructed a library of fourteen 18 nt UTR elements that generated a ∼38-fold range of protein output from a common T7 promoter. Application of this library to a three-gene β-carotene biosynthesis pathway enabled ∼20-fold variation in product titer through UTR assignment alone, without altering promoter identity or gene copy number. Together, these results establish a modular T7 RNAP expression framework in which cassette dosage defines overall transcriptional capacity while interchangeable UTR elements enable gene-specific tuning of expression output, providing a practical strategy for multigene expression control in yeast.

## Introduction

Predictable and independent control of multiple genes is a central challenge in synthetic biology, metabolic engineering, and genetic circuit design. In multigene systems, pathway performance often depends not only on the expression level of individual genes but also on their relative expression balance, making precise control of gene output essential for achieving desired cellular functions.^1^ However, co-expressed genetic modules compete for shared transcriptional and translational resources within the host cell, creating unintended interactions that can compromise expression predictability, circuit stability, and pathway performance.^2, 3^ These challenges have motivated the development of orthogonal expression systems that operate independently of endogenous regulatory networks and provide more predictable control over engineered genetic programs.^4^

*Saccharomyces cerevisiae* is a widely used host for synthetic biology and metabolic engineering owing to its well-characterized genetics, ease of genome engineering, and compatibility with complex eukaryotic protein expression.^5^ However, most engineered expression systems in yeast ultimately rely on the endogenous RNA polymerase II (RNAPII) machinery and its associated transcription factors, limiting the degree of transcriptional orthogonality that can be achieved.^6–8^ In multigene applications, the repeated use of a finite set of RNAPII-dependent promoters can further constrain circuit design and complicate predictable control of individual gene expression levels.

The bacteriophage-derived T7 RNA polymerase (T7 RNAP) offers an attractive alternative because it functions independently of host transcription factors and recognizes a highly specific 17 bp promoter sequence.^9–11^ This combination of transcriptional orthogonality, promoter specificity, and compact regulatory architecture has enabled widespread use of T7 RNAP-based expression systems in bacterial hosts, including *Escherichia coli*, *Bacillus megaterium*, and *Corynebacterium glutamicum*.^12, 13^ In particular, the small size and sequence specificity of the T7 promoter facilitate assembly of multigene constructs while maintaining a common transcriptional framework across multiple expression cassettes.^14^

Although T7 RNAP was first shown to initiate transcription in the yeast nucleus nearly four decades ago,^15^ productive expression from T7-derived transcripts has remained a longstanding challenge. Unlike transcripts generated by the endogenous RNAPII machinery, T7 RNAP transcripts do not naturally acquire the 5′ cap and 3′ poly(A) tail required for efficient mRNA maturation and utilization in eukaryotic cells,^16, 17^ Therefore, deficiencies in transcript stability, nuclear export, and downstream protein production have historically limited the practical application of T7 RNAP-based expression systems in yeast.

Recent advances have substantially improved the viability of this approach. Strategies including fusion of T7 RNAP to capping enzymes and subsequent optimization through directed evolution and machine learning have enabled efficient capping and productive expression from T7 promoter-driven transcripts in *S. cerevisiae*.^18–20^ These developments establish T7 RNAP as a practical orthogonal transcription platform in yeast and shift the remaining challenge from achieving expression itself toward controlling expression across multiple genes in a predictable and programmable manner.

Despite these advances, an important design challenge remains unresolved: how to achieve independent and predictable control of multiple genes within a shared T7 RNAP expression framework. Recent studies have demonstrated that T7 promoter variants with different transcriptional activities can be used to generate differential expression outputs across multiple cassettes.^19^. However, this strategy operates primarily at the level of transcription and relies on diversification of the promoter itself. As multigene constructs become larger and more complex, repeated use of promoter-derived regulatory elements can increase sequence redundancy and reduce design flexibility. More fundamentally, promoter-level tuning alone does not provide a modular mechanism for adjusting the output of individual genes while maintaining a common transcriptional architecture. A complementary regulatory layer capable of gene-specific tuning would therefore substantially expand the design space available for T7 RNAP-based multigene expression systems.

We reasoned that the 5′ untranslated region (UTR) could serve as such a regulatory layer. Short sequence elements within the 5’ UTR are known to influence protein output through post-transcriptional mechanisms, including effects on translation initiation and mRNA utilization.^21^ Unlike promoter engineering, which directly alters transcriptional input, diversification of the 5’ UTR offers a means to modulate gene-specific expression output while maintaining a common transcriptional driver. Moreover, because UTR elements are compact and readily interchangeable, they are well suited for modular assembly of multigene constructs. Diversification of the UTR region also increases sequence heterogeneity between adjacent expression cassettes, mitigating the risk of homologous recombination between repeated regulatory elements, which has been shown to cause structural instability in multigene yeast constructs.^22^ Together, these features suggest that the 5’ UTR can function as a compact and modular regulatory layer for gene-specific expression control within a shared T7 RNAP transcriptional framework.

Here, we report a modular design framework for T7 RNAP-based multigene expression in *S. cerevisiae* (Fig. 1A). We first established a pYTK-compatible expression architecture comprising interchangeable T7 promoter, UTR, polyadenylation signal, and terminator parts. Using a strain carrying a chromosomally integrated capping enzyme-fused T7 RNAP, we then examined the relationship between cassette dosage, transcript abundance, and expression output. To enable gene-specific control within a shared transcriptional framework, we developed a small library of fourteen interchangeable 18 nt UTR elements and demonstrated that UTR identity modulates protein output independently of major changes in transcript abundance. Finally, we applied this framework to a three-gene β-carotene biosynthesis pathway and showed that UTR assignment alone, without altering promoter identity or gene copy number, enables substantial tuning of pathway output. Together, these results establish a modular expression architecture in which cassette dosage determines overall transcriptional capacity while interchangeable UTR elements provide gene-specific control of protein output, offering a scalable strategy for multigene pathway engineering in eukaryotic systems.

**Figure 1.**
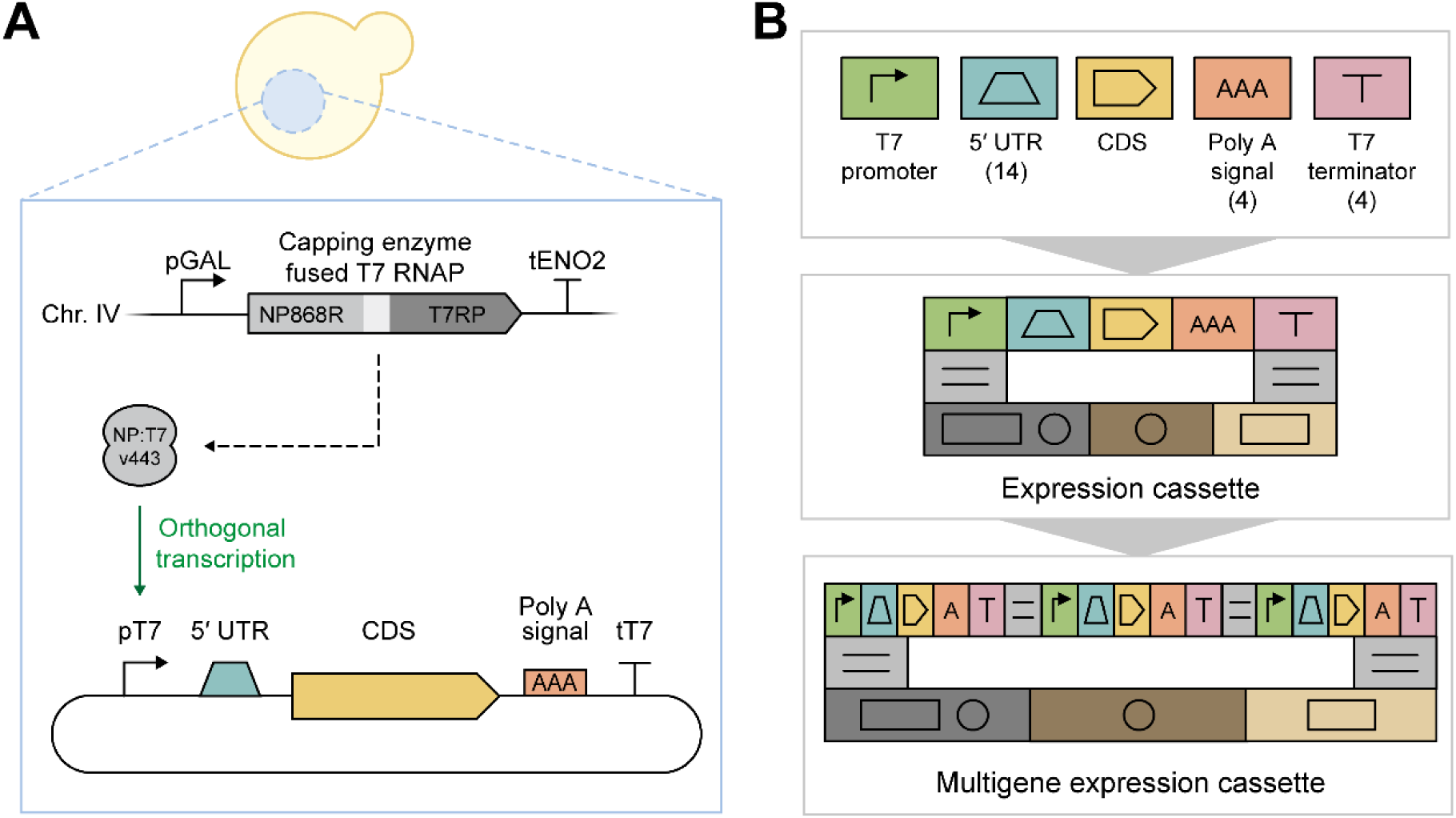
Construction of a modular pYTK-compatible T7 RNAP expression architecture in *S. cerevisiae.* (A) Overview of the orthogonal T7 RNAP expression system in *S. cerevisiae*. A capping enzyme-fused T7 RNAP (NP:T7 v443) was chromosomally integrated and used to drive transcription from T7 promoter-containing expression cassettes. (B) Modular organization of the pYTK-compatible T7 expression toolkit. T7 promoter, 5′ UTR, coding sequence (CDS), polyadenylation signal, and T7 terminator parts were assembled into single-gene or multigene expression cassettes using the pYTK framework.

## Results

### Construction of a modular pYTK-compatible T7 RNAP expression architecture

To establish a modular T7 RNAP-based expression platform in *S. cerevisiae*, we designed a set of pYTK-compatible bioparts for standardized and combinatorial cassette assembly. The T7 promoter part was engineered to incorporate features supporting both T7 RNAP-mediated transcription and downstream gene expression. Specifically, a 4 bp AT-rich sequence (AATT) was inserted immediately upstream of the promoter to enhance polymerase-promoter interactions,^23^ a 17 bp spacer sequence was incorporated downstream of the transcription start site to provide sufficient space for interchangeable 5′ UTR elements,^19^ and a yeast Kozak consensus sequence (AAAAAA) was positioned immediately upstream of the ATG start codon to support translation initiation in *S. cerevisiae* (Fig S1).^24, 25^ Incorporation of these elements required modification of the native YTK Type 2–Type 3 junction overhang from TATG to AATG.^26^ In addition to the promoter part, four T7 terminator variants^27^ (tT7 WT, tT7 hyb1, tT7 hyb2, tT7 hyb10) and four polyadenylation signal sequences^19^ (SV40, ADH1, SSA1, TDH1) were formatted as independent pYTK parts. Together, these components established a modular T7 RNAP expression architecture in which promoter, UTR, coding sequence, polyadenylation signal, and terminator elements could be independently assembled and exchanged within the pYTK framework (Fig. 1B; Table S2). To validate the redesigned pT7 part, we compared the previously reported pRS426-based construct lacking the AATT and Kozak elements^19^ with the pYTK-compatible version incorporating these modifications (Fig. 2A). The optimized construct exhibited a ∼1.8-fold increase in reporter expression relative to the original design (Fig. 2B), confirming the functionality of the redesigned promoter architecture within the pYTK framework. This optimized pT7 part was adopted as the standard transcriptional module for all subsequent experiments.

**Figure 2.**
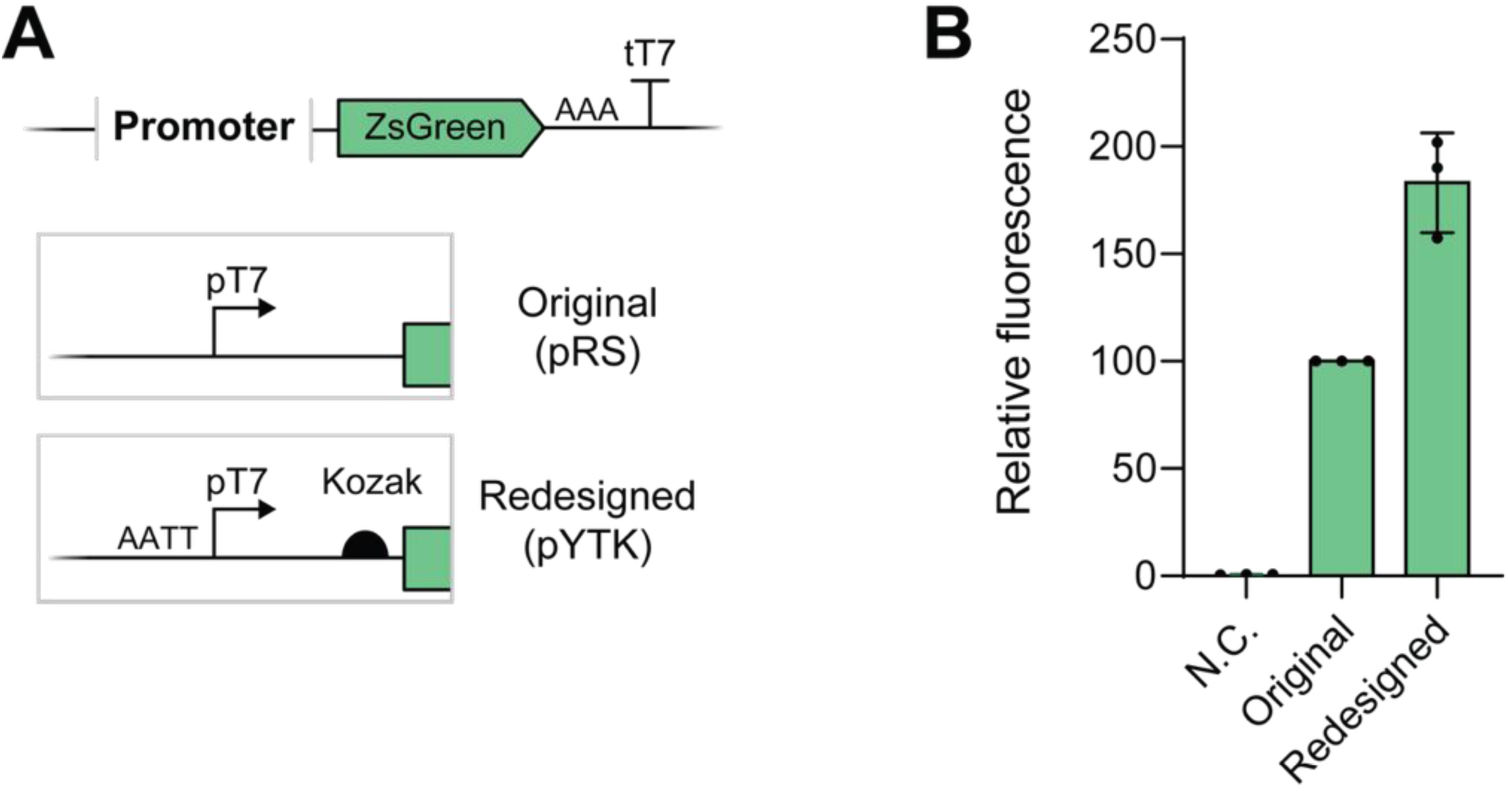
Redesign and validation of the pYTK-compatible T7 promoter part. (A) Schematic comparison of the original pRS-based T7 promoter configuration and the redesigned pYTK-compatible architecture containing an upstream AATT sequence and Kozak element. (B) Relative fluorescence of ZsGreen reporter constructs carrying the original or redesigned promoter architecture. Fluorescence values were normalized to the original construct. Data represent mean ± SD from three biological replicates. Statistical significance was determined by one-way ANOVA followed by Tukey’s multiple comparison test (p = 0.0129).

### Cassette dosage determines transcript abundance and expression output

To examine the effect of cassette dosage on expression output in the pYTK-compatible T7 RNAP system, we compared three expression configurations: single genomic integration (single copy), a centromeric plasmid (CEN/ARS; low copy), and a 2μ episomal plasmid (high copy) (Fig. 3A). Transcript abundance was quantified by RT-qPCR using *TAF10* as a housekeeping reference,^28^ and protein output was measured by flow cytometry. Relative to the high-copy condition, transcript levels reached only 4% and 8% in the single-copy and low-copy configurations, respectively, while protein output followed a parallel trend at 2% and 4% (Fig. 3B, C). Notably, the high-copy configuration produced transcript levels comparable to those obtained from the strong galactose-inducible *GAL1* promoter (∼95% of *pGAL*; Fig. S2), demonstrating that T7 RNAP-mediated transcription can achieve expression levels within the upper range of heterologous gene expression in yeast when present at sufficient dosage. Together, these results indicate that cassette dosage is a primary determinant of transcript abundance and overall expression output in the T7 RNAP system. Based on these findings, the high-copy 2μ plasmid format was adopted for all subsequent experiments.

**Figure 3.**
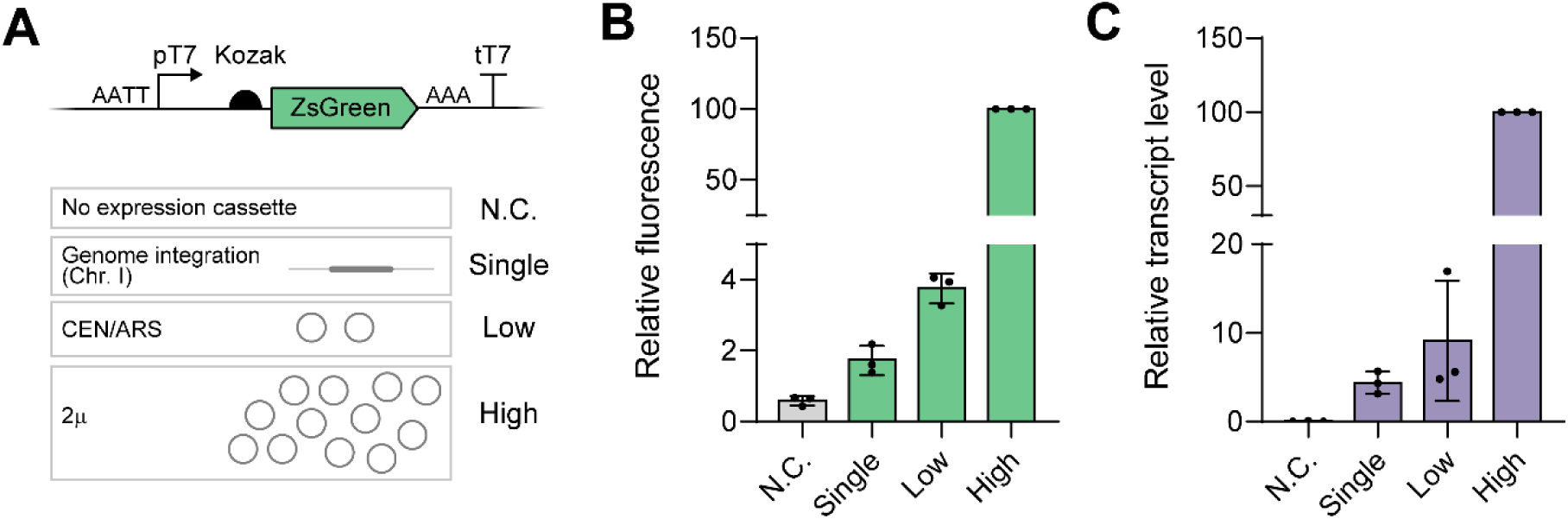
Cassette dosage determines transcript abundance and expression output in the T7 RNAP system. (A) Experimental design comparing expression cassette dosage using genomic integration (single copy), CEN/ARS plasmids (low copy), and 2μ plasmids (high copy). A strain lacking an expression cassette was included as a negative control (N.C.). (B) Relative ZsGreen fluorescence under different cassette dosage conditions. Fluorescence values were normalized to the high-copy condition. (C) Relative transcript abundance measured by RT-qPCR and normalized to the high-copy condition. *TAF10* was used as the reference gene. Data represent mean ± SD from three biological replicates. Statistical significance was assessed by one-way ANOVA followed by Tukey’s multiple comparison test (fluorescence, p < 0.0001; transcript abundance, p < 0.0001).

### UTR diversification enables gene-specific control of protein output

To enable gene-specific modulation of expression output within a shared T7 transcriptional framework, we separated the 5′ UTR from the pT7 part and established it as an independent pYTK component. Because one objective of the T7 RNAP architecture was to maintain compact expression cassettes suitable for multigene assembly, we focused on short 18 nt UTR elements. A small library of 14 such sequences, assembled from nucleotide motifs previously associated with translation initiation regulation,^21^ was cloned into a modular pYTK format together with a downstream Kozak consensus sequence. Each construct contained an identical T7 promoter and ZsGreen reporter and differed only in the 5′ UTR sequence. Expression levels were normalized to the pT7 promoter part described above, which served as the base construct (Fig. 4A, Base). Flow cytometric analysis revealed substantial variation in protein output across the 14 UTR variants, spanning an approximately 38-fold dynamic range relative to the base construct (Fig. 4B). Despite sharing an identical T7 promoter and coding sequence, individual UTR elements produced markedly different protein outputs, demonstrating that short sequence variations within the UTR region can strongly influence expression output. Based on their relative fluorescence values, the UTRs were classified into three expression tiers: High (UTR-A to UTR-C; 1.15–1.90× base), Mid (UTR-D to UTR-J; 0.52–0.93× base), and Low (UTR-K to UTR-N; 0.05–0.33× base) (Table S3). This tiered organization provided a practical framework for subsequent evaluation of gene-specific expression tuning in multigene pathway applications. To determine whether the observed differences in protein output arose from transcriptional or post-transcriptional effects, ZsGreen transcript abundance was quantified for the three High-tier UTR variants by RT-qPCR. Transcript levels for UTR-A, UTR-B, and UTR-C remained comparable to the base construct (mean relative transcript abundance of 1.09, 1.00, and 0.95, respectively), despite a 1.5-fold range in protein output across the same variants (1.15–1.90×; Fig. 4C). These results indicate that, at least within the High tier, differences in protein output are not explained by corresponding changes in transcript abundance. Instead, the UTR functions as a gene-specific regulatory element that modulates protein output while operating under a shared T7 transcriptional framework.

**Figure 4.**
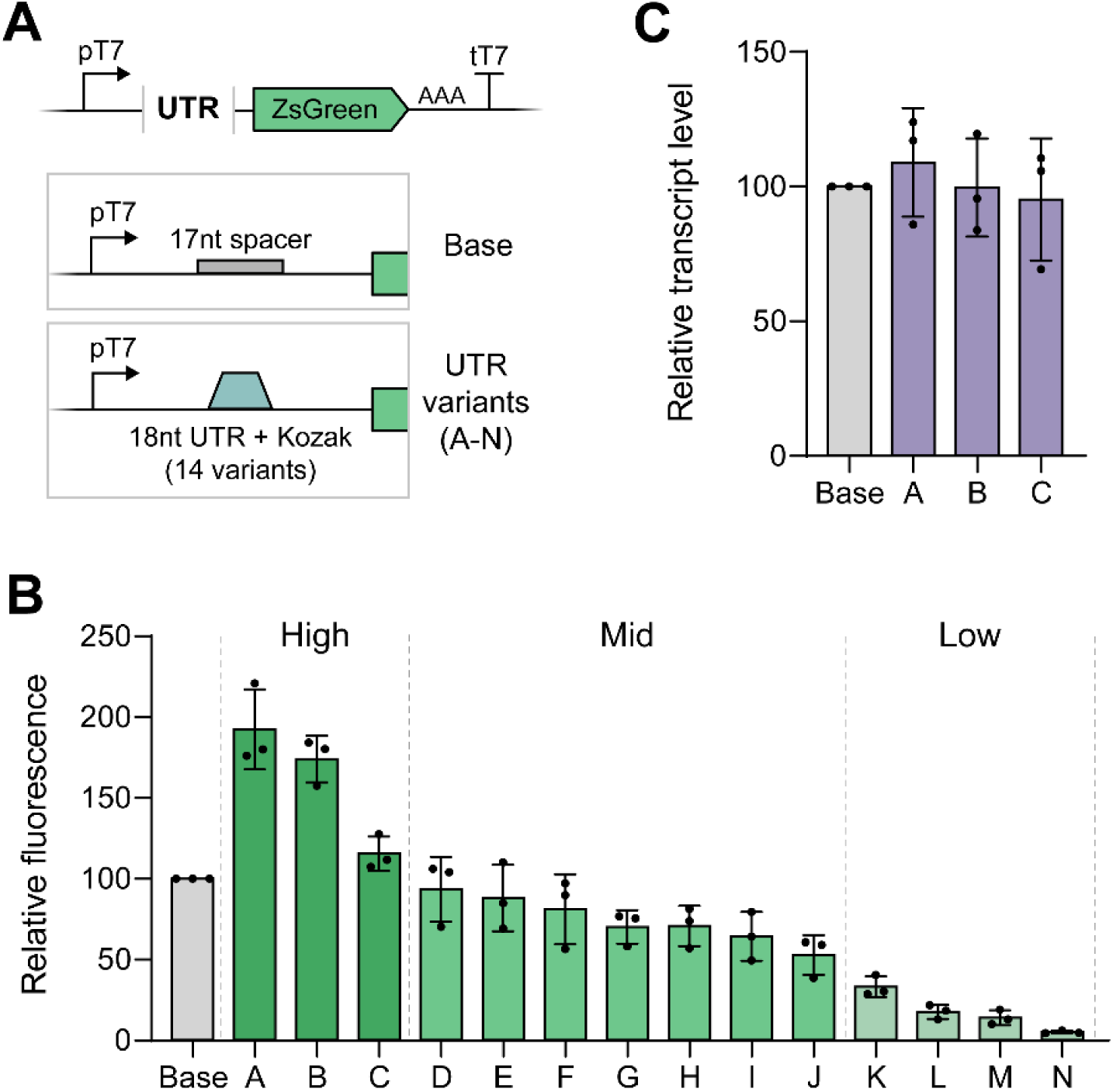
UTR diversification enables gene-specific tuning of protein output. (A) Design of the UTR library. The base construct contained the original 17 bp spacer sequence, whereas UTR variants contained one of fourteen 18 nt UTR elements and Kozak sequence positioned between the T7 promoter and the ZsGreen coding sequence. (B) Relative fluorescence of ZsGreen reporter constructs carrying different UTR variants. UTRs were grouped into High (A–C), Mid (D–J), and Low (K–N) expression tiers according to reporter output. Fluorescence values were normalized to the base construct. (C) Relative transcript abundance of the base construct and High-tier UTR variants measured by RT-qPCR. Transcript levels were normalized to the base construct using *TAF10* as the reference gene. Data represent mean ± SD from three biological replicates. Statistical significance was determined by one-way ANOVA followed by Tukey’s multiple comparison test (fluorescence, p < 0.0001; transcript abundance, p = 0.9826).

### Gene-specific UTR assignment enables pathway balancing within a shared transcriptional framework

To evaluate whether gene-specific UTR assignment could be used to tune expression across multiple genes, we assembled a three-gene β-carotene biosynthesis pathway consisting of CrtYB, CrtI, and CrtE. Each gene was expressed from an identical T7 promoter on a shared high-copy plasmid but assigned an independently selected UTR from the tiered library (Fig. 5A). Three representative UTRs per tier (High: UTR-A, –B, –C; Mid: UTR-G, –H, –I; Low: UTR-L, –M, –N) were tested in all tier combinations, yielding twenty-seven pathway strains with β-carotene titers ranging from 1.00 mg/L (Low–Low–Low) to 19.91 mg/L (High–High–High), representing an approximately 20-fold dynamic range in pathway output (Fig. 5B). The sum of individual UTR expression values showed a significant positive correlation with β-carotene titer (Spearman r = 0.70, p < 0.001, n = 27; Fig. 5C), indicating that combined expression potential is a primary determinant of pathway output. To determine whether pathway performance depended solely on overall expression strength or also on the distribution of expression among pathway enzymes, we constructed all six permutations of the three High-tier UTRs (UTR-A, UTR-B, and UTR-C) across CrtYB, CrtI, and CrtE. Despite containing the same set of UTR elements and therefore comparable overall expression potential, the six strains produced β-carotene titers ranging from 5.60 to 20.60 mg/L (Fig. 5D). These results demonstrate that gene-specific expression balancing can be achieved through UTR assignment alone, without altering promoter identity within the T7 RNAP architecture. Together, these findings establish UTR assignment as a gene-specific tuning parameter that enables implementation of pathway balancing within a shared T7 transcriptional framework, without requiring promoter diversification or copy-number optimization.

**Figure 5.**
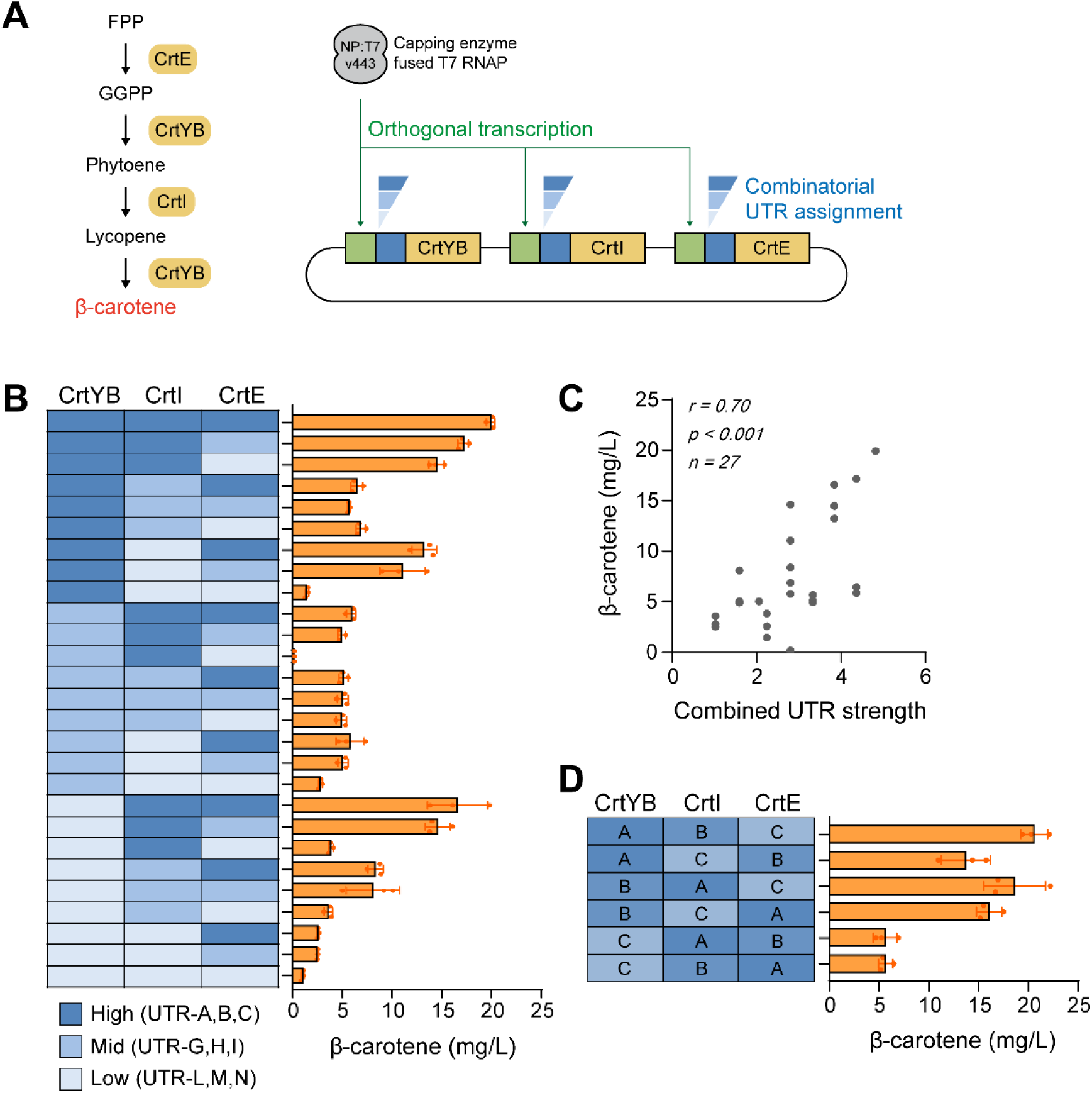
Gene-specific UTR assignment tunes β-carotene production in a multigene pathway. (A) Schematic of the β-carotene biosynthetic pathway and multigene T7 expression cassette. CrtYB, CrtI, and CrtE were expressed from independent transcription units sharing a common T7 transcriptional framework, while UTR identity was varied combinatorially. (B) β-Carotene production from pathway strains carrying different UTR combinations. Heatmap colors indicate High (UTR-A, UTR-B, UTR-C), Mid (UTR-G, UTR-H, UTR-I), and Low (UTR-L, UTR-M, UTR-N) UTR tiers. β-Carotene titers were quantified by HPLC. (C) Correlation between combined UTR strength and β-carotene production across all pathway strains (n = 27). Combined UTR strength was calculated as the sum of reporter-derived expression values associated with the assigned UTRs. (D) β-Carotene production from all six permutations of the three High-tier UTRs (UTR-A, UTR-B, and UTR-C) assigned across CrtYB, CrtI, and CrtE. Data represent mean ± SD from three biological replicates. Statistical significance was determined by one-way ANOVA followed by Tukey’s multiple comparison test (panels B and D, p < 0.0001). Correlation analysis in panel C was performed using Spearman’s rank correlation (r = 0.70, p < 0.001).

## Discussion

This study establishes a modular expression architecture for T7 RNAP-based multigene expression in *S. cerevisiae* by separating transcriptional and gene-specific expression control into distinct design layers. Previous efforts to improve T7 RNAP-mediated expression in yeast have focused primarily on enhancing transcript production or translational competence through engineering of the polymerase, capping machinery, or promoter sequence.^19, 20^ While these approaches have improved the performance of individual expression cassettes, strategies for independently controlling the output of multiple genes within a shared T7 transcriptional framework have remained limited. In contrast, the approach presented here maintains a common T7 promoter while introducing interchangeable UTR elements as independent regulatory parts, enabling differential control of individual genes without promoter diversification. By physically separating promoter and UTR functions within the pYTK framework, this architecture provides a scalable foundation for orthogonal multigene expression and establishes a shared transcriptional layer coupled to a gene-specific post-transcriptional layer.

The copy number and UTR experiments together reveal a hierarchical organization of expression control within the T7 RNAP system. High-copy plasmids were required to achieve transcript levels comparable to those obtained from the *GAL1* promoter, whereas low-copy and single-copy configurations produced substantially lower outputs. These results indicate that cassette dosage primarily determines the overall transcriptional capacity of the system. In contrast, diversification of the UTR sequence generated a broad range of protein outputs while maintaining comparable transcript levels among the tested high-expression variants, indicating that UTR identity influences gene-specific protein output through a post-transcriptional mechanism. This hierarchy establishes a modular design framework in which cassette dosage defines overall transcriptional capacity, while interchangeable UTR elements provide gene-specific control of protein output, enabling gene-specific tuning within a shared T7 transcriptional architecture without promoter diversification.

The β-carotene pathway experiments further demonstrate the practical utility of this architecture for multigene pathway balancing. Across the pathway strains, UTR assignment alone generated an approximately 20-fold range in β-carotene titers despite the use of identical T7 promoters and a common high-copy plasmid backbone. The significant correlation between combined UTR strength and product titer suggests that overall pathway output can be systematically modulated through selection of UTR combinations within a shared T7 transcriptional framework. While pathway strains with similar combined UTR strengths frequently exhibited substantial differences in β-carotene production, this observation is consistent with the well-established principle that pathway performance depends on the relative expression balance among constituent enzymes.^1^ The High-tier permutation experiments demonstrate that such gene-specific balancing can be achieved through UTR assignment alone, without altering promoter identity or gene copy number. Together, these results establish UTR assignment as a practical tuning strategy for implementing pathway balancing within a T7 RNAP-based orthogonal expression system. Rather than relying on promoter diversification to generate differential gene expression, this framework shifts pathway optimization to a compact and modular UTR layer while maintaining a common transcriptional architecture.

Several limitations of the current study should be considered. First, although the UTR library generated substantial differences in protein output, the molecular basis of these effects remains unresolved. The comparable transcript levels observed among the High-tier variants suggest a predominantly post-transcriptional mode of regulation, but whether the observed differences arise from altered translation initiation, mRNA stability, nuclear export, or a combination of these factors remains unclear. Moreover, RT-qPCR analysis was limited to the High-tier UTRs, and it therefore remains to be determined whether Mid– and Low-tier variants operate through similar mechanisms. A second limitation is that effective operation of the current architecture remained strongly dependent on the high-copy 2μ plasmid format, whereas low-copy and single-copy configurations produced substantially lower outputs. Strong copy number-dependent expression has also been observed in the pYTK system, where chromosomal integration, CEN/ARS plasmids, and 2μ plasmids produced distinct expression regimes.^26^ This dosage dependence likely reflects, at least in part, differences in template availability: in BY4741-derived haploid strains, CEN6/ARS4 plasmids have been reported at approximately 2–5 copies per cell, whereas auxotrophic-marker 2μ plasmids ranged from 14 to 34 copies per cell.^29^ Accordingly, the higher transcript abundance and protein output observed from the 2μ configuration in our system are consistent with increased availability of T7 promoter-containing templates. Nevertheless, further optimization will be required to improve expression from low-copy or genomic contexts.

Several opportunities exist to further expand the capabilities of the framework presented here. First, the current 14-element UTR library represents only a limited sampling of the sequence space available for post-transcriptional regulation. Expansion through rational design, high-throughput screening, or machine learning-guided sequence generation could increase both the dynamic range and resolution of gene-specific expression control while providing deeper insight into the sequence features that govern protein output. Second, the modular architecture of the pYTK framework enables straightforward incorporation of additional regulatory components. Future studies could combine the UTR library with T7 promoter variants or other regulatory elements to generate multidimensional expression landscapes spanning both transcriptional and post-transcriptional control. Finally, although this study was performed in *S. cerevisiae*, the design principles described here may be transferable to other eukaryotic hosts in which T7 RNAP-based expression systems have been established. Together, these extensions could broaden the utility of the platform as a general framework for orthogonal multigene expression and pathway engineering across diverse eukaryotic chassis.

In summary, this work demonstrates that T7 RNAP-based multigene expression in *S. cerevisiae* can be systematically organized into two separable control layers: cassette dosage, which sets overall transcriptional capacity, and UTR identity, which determines gene-specific protein output within a shared transcriptional framework. By establishing these layers within the pYTK modular cloning system, the architecture presented here provides a practical and scalable foundation for orthogonal multigene expression control in yeast. More broadly, the design principles described here—separating global transcriptional capacity from gene-specific post-transcriptional tuning—may provide a useful framework for programmable multigene expression in other eukaryotic systems where T7 RNAP-based platforms have been established.

## Methods

### Strains and Culture Conditions

*Saccharomyces cerevisiae* BY4742 (MATα *his3*Δ1 *leu2*Δ0 *lys2*Δ0 *ura3*Δ0) was used as the parental strain.^30^ A chromosomally integrated T7 RNAP expression cassette (pGal1-NPT7 v443-tENO2, carrying *HIS3* as a selectable marker) was introduced into the HO locus of BY4742 by homologous recombination, yielding the host strain used throughout this study (referred to hereafter as the T7 host strain).^19^ All reporter and pathway plasmids were transformed into this T7 host strain unless otherwise stated. Strains and plasmids used in this study are listed in Table S1. Yeast strains were cultivated at 30 °C in YPD or synthetic complete (SC) dropout medium with 2% glucose. Overnight cultures were initiated from glycerol stocks and grown for 20 h prior to induction. For induction, cultures were diluted 1:10 into fresh SC dropout medium containing 5% galactose and 2% raffinose and incubated at 30 °C for 18 h with orbital shaking. All expression measurements were performed at the 18 h time point.

### Yeast Transformation

Yeast transformation was performed using a modified lithium acetate/polyethylene glycol method.^31^ For chromosomal integration, plasmids were linearized by NotI digestion prior to transformation to promote integration by homologous recombination.

### Plasmid Construction

All plasmids were constructed using the yeast toolkit (pYTK) modular cloning framework.^26^ Parts not available in this library were synthesized as gene fragments (Twist Bioscience). Golden Gate assemblies were performed using BsmBI-v2 or BsaI-HF v2 according to standard pYTK procedures.^26^ Thermocycler conditions: 25 cycles of 42 °C for 1 min 30 s and 16 °C for 3 min, followed by 60 °C for 10 min and 80 °C for 10 min. All reporter constructs were based on a pT7–ZsGreen–poly(A)–tT7 architecture. High-copy and low-copy plasmids were constructed using 2μ and CEN/ARS origins, respectively. Single-copy constructs were integrated into the IntI genomic locus using the MYT integration platform^32^. Carotenoid biosynthesis genes (*crtYB*, *crtI*, *crtE*) from *Xanthophyllomyces dendrorhous* were synthesized as codon-optimized gene fragments based on previously reported sequences.^33, 34^ Four T7 terminator variants^27^ (tT7 WT, tT7 hyb1, tT7 hyb2, tT7 hyb10) and four polyadenylation signal sequences^19^ (SV40, ADH1, SSA1, TDH1) were formatted as independent pYTK parts. A library of 14 synthetic 5′ UTR elements (18 nt each) was assembled from short nucleotide sequences previously reported to influence translation initiation.^21^ Each element was cloned immediately downstream of the T7 promoter together with a downstream Kozak consensus sequence (AAAAAA) immediately upstream of the ATG start codon and formatted as an independent pYTK part. Each element was designated UTR-A through UTR-N in descending order of protein expression output, as determined by flow cytometry screening described below. Sequences and corresponding expression data for all 14 elements are provided in Table S3.

### *Escherichia coli* Transformation and Plasmid Verification

*Escherichia coli* DH5α was used for plasmid propagation. Chemically competent cells were prepared by the Inoue method^35^ and selected on LB agar with appropriate antibiotics. All plasmid inserts were verified by Sanger sequencing (Macrogen, Seoul).

### Flow Cytometry

Cultures were harvested 18 h post-induction and diluted 1:100 in phosphate-buffered saline (PBS). Flow cytometry was performed on a CytoFLEX flow cytometer (Beckman Coulter). ZsGreen fluorescence was measured using 488 nm excitation and a 525/40 nm bandpass emission filter. A minimum of 10,000 gated events were collected per sample, and mean fluorescence intensity (MFI) was recorded. Data were analyzed using FlowJo software (BD Biosciences).

### RT-qPCR Analysis

Total RNA was extracted from cultures harvested 18 h post-induction using the RNeasy Mini Kit (Qiagen) according to the manufacturer’s instructions. RNA concentration and purity were assessed by NanoDrop spectrophotometry. First-strand cDNA synthesis was performed using ReverTra Ace qPCR RT Master Mix (TOYOBO). Quantitative PCR was conducted using BioFACT 2× Real-Time PCR Master Mix with SFCgreen I on a CFX96 Real-Time PCR Detection System (Bio-Rad). Data were analyzed using Bio-Rad CFX Manager software (v3.1). Transcript levels were quantified by the ΔΔCt method using *TAF10* as a constitutively expressed reference gene.^28^ Relative transcript abundance was calculated using the 2^−ΔΔCt^ method.^36^ The reference condition used for normalization is indicated in each figure. Primer sequences are listed in Table S5.

### β-Carotene Quantification

β-Carotene extraction and quantification were performed as previously described,^34^ with minor modifications. β-Carotene production was quantified from cultures harvested 18 h after galactose induction. For each sample, 1 mL of culture was collected and lyophilized. Lyophilized pellets were resuspended in 800 μL dimethyl sulfoxide (DMSO) and incubated at room temperature for 1 h with agitation. Hexane (800 μL) was added, and samples were vortexed thoroughly. After centrifugation, the upper organic phase was transferred to a fresh tube and evaporated at 40 °C. Dried residues were dissolved in 100 μL acetone for HPLC analysis. HPLC analysis was performed on an Agilent 1200 system equipped with a ZORBAX Eclipse Plus C18 column (4.6 × 150 mm, 3.5 μm). The mobile phase consisted of methanol/acetonitrile/dichloromethane (21:21:8, v/v/v) at an isocratic flow rate of 1.0 mL/min. Column temperature was maintained at 30 °C, and detection was performed at 450 nm. Quantification was based on peak area using a β-carotene external standard curve (Sigma-Aldrich).

### Statistical Analysis

All experiments were performed with three independent biological replicates. Data are presented as mean ± standard deviation (SD). Statistical significance between two groups was assessed by a two-tailed Student’s t-test. Comparisons among three or more groups were analyzed by one-way ANOVA followed by Tukey’s multiple comparison test. Rank correlation between combined UTR strength and β-carotene titer was assessed using Spearman’s rank correlation coefficient. Differences were considered statistically significant at p < 0.05. All statistical analyses and graph generation were performed using GraphPad Prism (v11.0.1).

## Author Contributions

E.J. and S.H. contributed equally to this work. E.J., S.H., S.-G.W., and D.-H.L. conceived the study. E.J. and S.H. performed the experiments. E.J., S.H., and S.-G.W. analyzed the data. E.J., S.H., and D.-H.L. wrote the paper. D.-H.L. supervised the project and revised the manuscript. All authors reviewed and approved the final manuscript.

## Acknowledgments

We thank Andrew D. Ellington and members of the Ellington laboratory for providing the NP:T7v443 expression plasmid and reporter plasmids used in this study. We also thank members of the Synthetic Biology Research Center for helpful discussions and technical assistance.

## Data Availability

All data supporting the findings of this study are available within the article and its Supporting Information files.

