## Supplementary Information for "A Modular T7 RNAP Expression Architecture for Orthogonal Multigene Expression in Yeast"

### Supporting Figure

**Figure S1.** Sequence comparison of the original and redesigned T7 promoter modules.
**Figure S2.** Comparison of expression driven by the T7 RNAP system and the *GAL1* promoter.

### Supporting Tables

**Table S1.** Strains and plasmids used in this study.
**Table S2.** Sequences of T7 RNAP expression parts.
**Table S3.** UTR library sequences and expression characteristics.
**Table S4.** β-carotene pathway strain designs and production data.
**Table S5.** Primers used in this study.

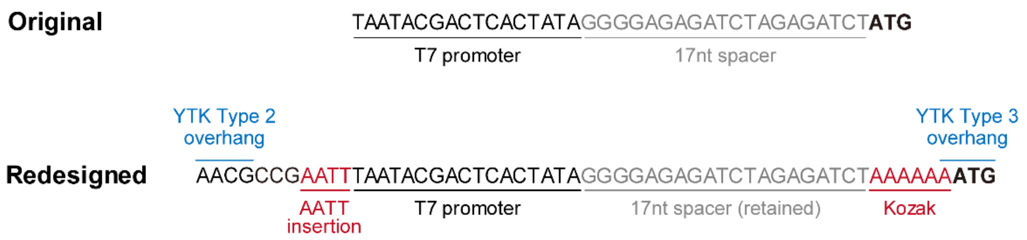


**Figure S1. Sequence architecture of the original and redesigned T7 promoter modules used in Figure 2.** The redesigned construct incorporates a 4-bp AATT insertion, a yeast Kozak consensus sequence (AAAAAA), and pYTK-compatible Type 2 and Type 3 assembly overhangs while retaining the original 17-nt spacer sequence between the T7 promoter and start codon. To accommodate the Kozak sequence, the standard pYTK Type 3 overhang (TATG) was modified to AATG.

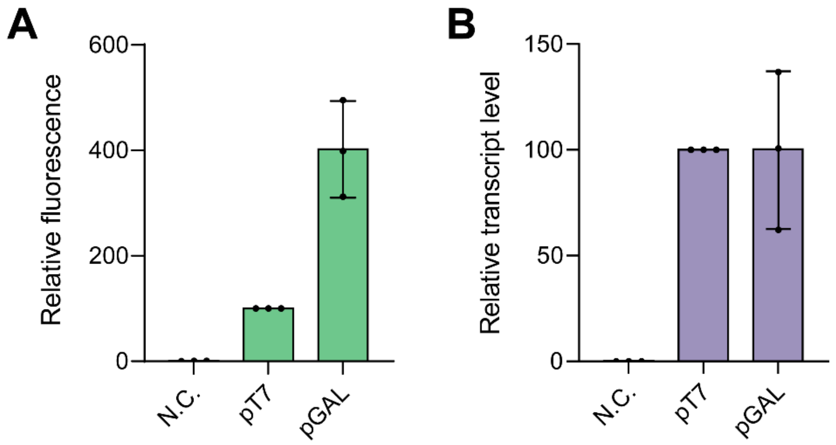


**Figure S2. Comparison of expression driven by the T7 RNAP system and the *GAL1* promoter.** (A) Relative fluorescence of ZsGreen expressed from the optimized T7 RNAP expression cassette (pT7) or the *GAL1* promoter (pGAL). Fluorescence values were normalized to the pT7 condition. N.C., negative control lacking an expression cassette. (B) Relative transcript levels of ZsGreen measured by RT-qPCR and normalized to the pT7 condition. Data represent the mean ± SD of three biological replicates.

**Table S1.** Strains and plasmids used in this study.

| **Plasmid** | | | | |
|---|---|---|---|---|
| **Category** | **Name** | **Description** | **Selection marker** | **Source** |
| T7 RNAP expression | ES443 | pGAL-NPT7v443-tENO2;<br>HIS3, HO homology arm | Kan | Kar et al. (2025) |
| Reporter | ES205 | Original pT7-ZsG-SV40-tT7-2-micron | Amp | This study |
| Reporter | pEJ001 | pYTK AT-pT7-IT-Kozak-ZsG | Amp | This study |
| Reporter | pEJ002 | pYTK pT7-ZsG-SV40-tT7-2-micron | Amp | This study |
| Reporter | pEJ003 | pYTK pT7-ZsG-SV40-tT7-CEN | Amp | This study |
| Reporter | pEJ004 | pMYT pT7-ZsG-SV40-tT7;<br>integration at Chr. I | Amp | This study |
| Reporter | pEJ005 | pYTK pGal_ZsG_tENO2 | Kan | This study |
| Reporter | pEJ006 | pYTK pT7-UTRa-ZsG-SV40-tT7 | Kan | This study |
| Reporter | pEJ007 | pYTK pT7-UTRb-ZsG-SV40-tT7 | Kan | This study |
| Reporter | pEJ008 | pYTK pT7-UTRc-ZsG-SV40-tT7 | Kan | This study |
| Reporter | pEJ009 | pYTK pT7-UTRd-ZsG-SV40-tT7 | Kan | This study |
| Reporter | pEJ010 | pYTK pT7-UTRe-ZsG-SV40-tT7 | Kan | This study |
| Reporter | pEJ011 | pYTK pT7-UTRf-ZsG-SV40-tT7 | Kan | This study |
| Reporter | pEJ012 | pYTK pT7-UTRg-ZsG-SV40-tT7 | Kan | This study |
| Reporter | pEJ013 | pYTK pT7-UTRh-ZsG-SV40-tT7 | Kan | This study |
| Reporter | pEJ014 | pYTK pT7-UTRi-ZsG-SV40-tT7 | Kan | This study |
| Reporter | pEJ015 | pYTK pT7-UTRj-ZsG-SV40-tT7 | Kan | This study |
| Reporter | pEJ016 | pYTK pT7-UTRk-ZsG-SV40-tT7 | Kan | This study |
| Reporter | pEJ017 | pYTK pT7-UTRl-ZsG-SV40-tT7 | Kan | This study |
| Reporter | pEJ018 | pYTK pT7-UTRm-ZsG-SV40-tT7 | Kan | This study |
| Reporter | pEJ019 | pYTK pT7-UTRn-ZsG-SV40-tT7 | Kan | This study |
| β-carotene pathway | pEJ_S01[a] | UTRb-CrtYB-UTRc-CrtI-UTRa-CrtE | Kan | This study |
| β-carotene pathway | pEJ_S02 | UTRb-CrtYB-UTRg-CrtI-UTRa-CrtE | Kan | This study |
| β-carotene pathway | pEJ_S03 | UTRb-CrtYB-UTRl-CrtI-UTRa-CrtE | Kan | This study |
| β-carotene pathway | pEJ_S04 | UTRg-CrtYB-UTRb-CrtI-UTRa-CrtE | Kan | This study |
| β-carotene pathway | pEJ_S05 | UTRg-CrtYB-UTRh-CrtI-UTRa-CrtE | Kan | This study |
| β-carotene pathway | pEJ_S06 | UTRg-CrtYB-UTRl-CrtI-UTRa-CrtE | Kan | This study |
| β-carotene pathway | pEJ_S07 | UTRl-CrtYB-UTRb-CrtI-UTRa-CrtE | Kan | This study |
| β-carotene pathway | pEJ_S08 | UTRl-CrtYB-UTRg-CrtI-UTRa-CrtE | Kan | This study |
| β-carotene pathway | pEJ_S09 | UTRl-CrtYB-UTRm-CrtI-UTRa-CrtE | Kan | This study |
| β-carotene pathway | pEJ_S10 | UTRa-CrtYB-UTRb-CrtI-UTRg-CrtE | Kan | This study |
| β-carotene pathway | pEJ_S11 | UTRa-CrtYB-UTRh-CrtI-UTRg-CrtE | Kan | This study |
| β-carotene pathway | pEJ_S12 | UTRa-CrtYB-UTRl-CrtI-UTRg-CrtE | Kan | This study |
| β-carotene pathway | pEJ_S13 | UTRh-CrtYB-UTRa-CrtI-UTRg-CrtE | Kan | This study |
| β-carotene pathway | pEJ_S14 | UTRh-CrtYB-UTRi-CrtI-UTRg-CrtE | Kan | This study |

| | | | | |
|---|---|---|---|---|
| β-carotene pathway | pEJ_S15 | UTRh-CrtYB-UTRl-CrtI-UTRg-CrtE | Kan | This study |
| β-carotene pathway | pEJ_S16 | UTRl-CrtYB-UTRa-CrtI-UTRg-CrtE | Kan | This study |
| β-carotene pathway | pEJ_S17 | UTRl-CrtYB-UTRh-CrtI-UTRg-CrtE | Kan | This study |
| β-carotene pathway | pEJ_S18 | UTRl-CrtYB-UTRm-CrtI-UTRg-CrtE | Kan | This study |
| β-carotene pathway | pEJ_S19 | UTRa-CrtYB-UTRb-CrtI-UTRl-CrtE | Kan | This study |
| β-carotene pathway | pEJ_S20 | UTRa-CrtYB-UTRg-CrtI-UTRl-CrtE | Kan | This study |
| β-carotene pathway | pEJ_S21 | UTRa-CrtYB-UTRm-CrtI-UTRl-CrtE | Kan | This study |
| β-carotene pathway | pEJ_S22 | UTRg-CrtYB-UTRa-CrtI-UTRl-CrtE | Kan | This study |
| β-carotene pathway | pEJ_S23 | UTRg-CrtYB-UTRh-CrtI-UTRl-CrtE | Kan | This study |
| β-carotene pathway | pEJ_S24 | UTRg-CrtYB-UTRm-CrtI-UTRl-CrtE | Kan | This study |
| β-carotene pathway | pEJ_S25 | UTRm-CrtYB-UTRa-CrtI-UTRl-CrtE | Kan | This study |
| β-carotene pathway | pEJ_S26 | UTRm-CrtYB-UTRg-CrtI-UTRl-CrtE | Kan | This study |
| β-carotene pathway | pEJ_S27 | UTRm-CrtYB-UTRn-CrtI-UTRl-CrtE | Kan | This study |
| β-carotene pathway | pEJ_H2 | UTRc-CrtYB-UTRb-CrtI-UTRa-CrtE | Kan | This study |
| β-carotene pathway | pEJ_H3 | UTRa-CrtYB-UTRc-CrtI-UTRb-CrtE | Kan | This study |
| β-carotene pathway | pEJ_H4 | UTRc-CrtYB-UTRa-CrtI-UTRb-CrtE | Kan | This study |
| β-carotene pathway | pEJ_H5 | UTRa-CrtYB-UTRb-CrtI-UTRc-CrtE | Kan | This study |
| β-carotene pathway | pEJ_H6 | UTRb-CrtYB-UTRa-CrtI-UTRc-CrtE | Kan | This study |
| **Yeast strain** | | | | |
| **Category** | **Name** | **Description** | **Selection marker** | **Source** |
| T7 host | yDL001 | BY4742; HO::pGAL1-NP:T7 v443-tENO2 | HIS3 | This study |
| Promoter comparison | yDL011 | yDL001; ES205 | URA3 | This study |
| Promoter comparison | yEJ001 | yDL001; pEJ001 | URA3 | This study |
| Dosage comparison | yEJ002 | yDL001; pEJ002 | URA3 | This study |
| Dosage comparison | yEJ003 | yDL001; pEJ003 | URA3 | This study |
| Dosage comparison | yEJ004 | yDL001; pEJ004 | URA3 | This study |
| Dosage comparison | yEJ005 | yDL001; pEJ005 | URA3 | This study |
| UTR variant | yEJ006 | yDL001; pEJ006 | LEU2 | This study |
| UTR variant | yEJ007 | yDL001; pEJ007 | LEU2 | This study |
| UTR variant | yEJ008 | yDL001; pEJ008 | LEU2 | This study |
| UTR variant | yEJ009 | yDL001; pEJ009 | LEU2 | This study |
| UTR variant | yEJ010 | yDL001; pEJ010 | LEU2 | This study |
| UTR variant | yEJ011 | yDL001; pEJ011 | LEU2 | This study |
| UTR variant | yEJ012 | yDL001; pEJ012 | LEU2 | This study |
| UTR variant | yEJ013 | yDL001; pEJ013 | LEU2 | This study |
| UTR variant | yEJ014 | yDL001; pEJ014 | LEU2 | This study |
| UTR variant | yEJ015 | yDL001; pEJ015 | LEU2 | This study |
| UTR variant | yEJ016 | yDL001; pEJ016 | LEU2 | This study |
| UTR variant | yEJ017 | yDL001; pEJ017 | LEU2 | This study |

| | | | | |
|---|---|---|---|---|
| UTR variant | yEJ018 | yDL001; pEJ018 | LEU2 | This study |
| UTR variant | yEJ019 | yDL001; pEJ019 | LEU2 | This study |
| β-carotene pathway | yEJ020[a] | yDL001; pEJ_S01 | LEU2 | This study |
| β-carotene pathway | yEJ021 | yDL001; pEJ_S02 | LEU2 | This study |
| β-carotene pathway | yEJ022 | yDL001; pEJ_S03 | LEU2 | This study |
| β-carotene pathway | yEJ023 | yDL001; pEJ_S04 | LEU2 | This study |
| β-carotene pathway | yEJ024 | yDL001; pEJ_S05 | LEU2 | This study |
| β-carotene pathway | yEJ025 | yDL001; pEJ_S06 | LEU2 | This study |
| β-carotene pathway | yEJ026 | yDL001; pEJ_S07 | LEU2 | This study |
| β-carotene pathway | yEJ027 | yDL001; pEJ_S08 | LEU2 | This study |
| β-carotene pathway | yEJ028 | yDL001; pEJ_S09 | LEU2 | This study |
| β-carotene pathway | yEJ029 | yDL001; pEJ_S10 | LEU2 | This study |
| β-carotene pathway | yEJ030 | yDL001; pEJ_S11 | LEU2 | This study |
| β-carotene pathway | yEJ031 | yDL001; pEJ_S12 | LEU2 | This study |
| β-carotene pathway | yEJ032 | yDL001; pEJ_S13 | LEU2 | This study |
| β-carotene pathway | yEJ033 | yDL001; pEJ_S14 | LEU2 | This study |
| β-carotene pathway | yEJ034 | yDL001; pEJ_S15 | LEU2 | This study |
| β-carotene pathway | yEJ035 | yDL001; pEJ_S16 | LEU2 | This study |
| β-carotene pathway | yEJ036 | yDL001; pEJ_S17 | LEU2 | This study |
| β-carotene pathway | yEJ037 | yDL001; pEJ_S18 | LEU2 | This study |
| β-carotene pathway | yEJ038 | yDL001; pEJ_S19 | LEU2 | This study |
| β-carotene pathway | yEJ039 | yDL001; pEJ_S20 | LEU2 | This study |
| β-carotene pathway | yEJ040 | yDL001; pEJ_S21 | LEU2 | This study |
| β-carotene pathway | yEJ041 | yDL001; pEJ_S22 | LEU2 | This study |
| β-carotene pathway | yEJ042 | yDL001; pEJ_S23 | LEU2 | This study |
| β-carotene pathway | yEJ043 | yDL001; pEJ_S24 | LEU2 | This study |
| β-carotene pathway | yEJ044 | yDL001; pEJ_S25 | LEU2 | This study |
| β-carotene pathway | yEJ045 | yDL001; pEJ_S26 | LEU2 | This study |
| β-carotene pathway | yEJ046 | yDL001; pEJ_S27 | LEU2 | This study |
| β-carotene pathway | yEJ048 | yDL001; pEJ_H2 | LEU2 | This study |
| β-carotene pathway | yEJ049 | yDL001; pEJ_H3 | LEU2 | This study |
| β-carotene pathway | yEJ050 | yDL001; pEJ_H4 | LEU2 | This study |
| β-carotene pathway | yEJ051 | yDL001; pEJ_H5 | LEU2 | This study |
| β-carotene pathway | yEJ052 | yDL001; pEJ_H6 | LEU2 | This study |

[a]pEJ_S01 served as both the S01 and H1 constructs in the β-carotene pathway experiments.

**Table S2.** Sequences of T7 RNAP expression parts.

| Part Type / Category | | Part Name | Length (including overhangs) | Sequence (5'→3', including overhangs) |
|---|---|---|---|---|
| 2a | Promoter[a] | pT7-AATT | 34 | AACGCCGAATTTAATACGACTCACTATAGGGAGA |
| 2b | 5'UTR[b] | UTR-A | 31 | GAGAGTAGGTCTACTACACAGTAAAAAAATG |
| 2b | 5'UTR[b] | UTR-B | 31 | GAGAAAGGGCTGCAAATATCTGAAAAAAATG |
| 2b | 5'UTR[b] | UTR-C | 31 | GAGAGGTGCGCGTGGAGCTTCTAAAAAAATG |
| 2b | 5'UTR[b] | UTR-D | 31 | GAGAAGGAGAGGCTGGTTGCTAAAAAAAATG |
| 2b | 5'UTR[b] | UTR-E | 31 | GAGAGTTAAGACCCGGGAATGCAAAAAAATG |
| 2b | 5'UTR[b] | UTR-F | 31 | GAGAGTCGCGTGCAGTCATCCTAAAAAAATG |
| 2b | 5'UTR[b] | UTR-G | 31 | GAGAGTTGGTTGGGTGAGCAGTAAAAAAATG |
| 2b | 5'UTR[b] | UTR-H | 31 | GAGAGGACAAGGGGGTGTTTGAAAAAAAATG |
| 2b | 5'UTR[b] | UTR-I | 31 | GAGAGGGCGGTGGACTGACATCAAAAAAATG |
| 2b | 5'UTR[b] | UTR-J | 31 | GAGAGGCCTGGGCGCGGCTGAGAAAAAAATG |
| 2b | 5'UTR[b] | UTR-K | 31 | GAGAGGGCAGCATCGCTGCGACAAAAAAATG |
| 2b | 5'UTR[b] | UTR-L | 31 | GAGAATTCCGGATGCCGGTTCGAAAAAAATG |
| 2b | 5'UTR[b] | UTR-M | 31 | GAGAGATGCTGGGGTTTTGCATAAAAAAATG |
| 2b | 5'UTR[b] | UTR-N | 31 | GAGAAGGGGGGCGGGGAGTGATAAAAAAATG |
| 3a' | CDS[c] | ZsGreen | 701 | AATGGCACAGAGTGAGCATGGACTTACGGAAGAGATGAC<br>GATGAAATATAGAATGGAGGGTTGTGTCGATGGACACAAG<br>TTTGTTATTACGGGGGAGGGAATCGGATACCCCTTTAAAG<br>GCAAACAGGCCATAAATTTATGTGTCGTTGAGGGCGGTC<br>CACTACCATTTGCTGAGGACATTTTATCAGCCGCTTTTATG<br>TACGGTAACAGGGTGTTCACCGAATACCCTCAGGACATC<br>GTCGACTACTTCAAAAATAGCTGCCCGGCCGGTTATACG<br>TGGGATAGAAGCTTTCTATTTGAGGATGGTGCCGTATGTAT<br>CTGCAACGCCGACATAACCGTATCCGTCGAAGAAAATTG<br>CATGTACCACGAATCCAAATTCTATGGTGTGAACTTTCCT<br>GCTGATGGTCCCGTTATGAAAAAAATGACGGATAACTGG<br>GAGCCAAGCTGCGAGAAGATCATCCCAGTGCCCAAGCA<br>AGGCATTCTTAAAGGTGACGTATCTATGTACCTACTGCTAA<br>AAGATGGAGGGAGACTAAGGTGCCAATTCGATACAGTTTA<br>TAAAGCCAAGTCAGTCCCGCGTAAGATGCCCGATTGGCA<br>CTTTATCCAACACAAGTTGACAAGGGAGGATAGAAGCGA<br>TGCAAAAAATCAGAAGTGGCATTTAACCGAGCACGCTAT<br>CGCTAGTGGCTCCGCGCTGCCTTAAATCC |
| 4a' | Poly A signal[d] | SV40 | 285 | ATCCCTTCGAGCAGACATGATAAGATACATTGATGAGTTT<br>GGACAAACCACAACTAGAATGCAGTGAAAAAAATGCTTTA<br>TTTGTGAAATTTGTGATGCTATTGCTTTATTTGTAACCATTAT<br>AAGCTGCAATAAACAAGTTAACAACAACAATTGCATTCATT<br>TTATGTTTCAGGTTCAGGGGGAGATGTGGGAGGTTTTTTAA<br>AGCAAGTAAAACCTCTACAAATGTGGTAAAATCCGATAGG<br>ATCCTAACTCGAGACTGAAAGCTTAATTAGCAGCATTGGC |
| 4a' | Poly A signal[d] | tADH1 | 233 | ATCCGCGAATTTCTTATGATTTATGATTTTTATTATTAAATAA<br>GTTATAAAAAAAATAAGTGTATACAAATTTTAAAGTGACTCT<br>TAGGTTTTAAAACGAAAATTCTTATTCTTGAGTAACTCTTTC<br>CTGTAGGTCAGGTTGCTTTCTCAGGTATAGCATGAGGTCG<br>CTCTTATTGACCACACCTCTACCGGCATGCCGAGCAAAT<br>GCCTGCAAATCGCTCCCCATTTCTGGC |
| 4a' | Poly A signal[d] | tSSA1 | 233 | ATCCGCCAATTGGTGCGGCAATTGATAATAACGAAAATGT<br>CTTTTAATGATCTGGGTATAATGAGGAATTTTCCGAACGTTT<br>TTACTTTATATATATATATACATGTAACATATATTCTATACGC<br>TATAGAGAAAGGAAATTTTTCAATTAAAAAAAAAAATAGAGAA<br>AGAGTTTCACTTCTTGATTATCGCTAACACTAATGGTTGAA<br>GTACTGCTACTTTAATTTTATTGGC |
| 4a' | Poly A signal[d] | tTDH1 | 232 | ATCCATAAAGCAATCTTGATGAGGATAATGATTTTTTTTTGA<br>ATATACATAAATACTACCGTTTTTCTGCTAGATTTTGTGATG |

| | | | | |
|---|---|---|---|---|
| | | | | ACGTAAATAAGTACATATTACTTTTTAAGCCAAGACAAGATT AAGCATTAACTTTACCCTTTTCTTTCTAAGTTTCAATATTAGT TATCACTGTTTAAAAGTTATGGCGAGAACGTCGGCGGTTA AAATATATTACCCTGAACGTGGC |
| 4b | Terminator[e] | tT7 WT | 55 | TGGCTAGCATAACCCCTTGGGGCCTCTAAACGGGTCTTG AGGGGTTTTTTGGCTG |
| 4b | Terminator[e] | tT7 hyb1 | 50 | TGGCAAACAGATAGGCCCTCTTCGGAGGGCCTATCTGTT TTTTTTTGCTG |
| 4b | Terminator[e] | tT7 hyb2 | 46 | TGGCAACAGATAGGCCTCTTCGGAGGCCTATCTGTTTTTT TTGCTG |
| 4b | Terminator[e] | tT7 hyb10 | 86 | TGGCAGATAACAGATACTTCGGTATCTGTTATCTGTTTTTTT TCAACAGATAGCCGCGTTCGCGCGGCTATCTGTTTTTTTT GCTG |

[a]Promoter part sequence derived from the canonical T7 promoter and modified to include an upstream AATT enhancer sequence and GGGAGA initial transcribed sequence.

[b]UTR sequences derived from Zhou et al. (2003); formatted as an independent pYTK part with a Kozak sequence.

[c]ZsGreen reporter sequence derived from Kar et al. (2025); formatted as an independent pYTK part with a modified Type 3 junction overhang (AATG).

[d]Polyadenylation sequences adapted from Kar et al. (2025).

[e]Terminator sequences adapted from Calvopina-Chavez et al. (2022)

**Table S3.** UTR library sequences and expression characteristics.

| UTR ID | Sequence (5'→3') | Relative fluorescence (base=1) | Tier |
|---|---|---|---|
| A | GTAGGTCTACTACACAGT | 1.92 | High |
| B | AAGGGCTGCAAATATCTG | 1.74 | High |
| C | GGTGCGCGTGGAGCTTCT | 1.16 | High |
| D | AGGAGAGGCTGGTTGCTA | 0.94 | Mid |
| E | GTTAAGACCCGGGAATGC | 0.88 | Mid |
| F | GTCGCGTGCAGTCATCCT | 0.81 | Mid |
| G | GTTGGTTGGGTGAGCAGT | 0.70 | Mid |
| H | GGACAAGGGGGTGTTTGA | 0.71 | Mid |
| I | GGGCGGTGGACTGACATC | 0.64 | Mid |
| J | GGCCTGGGCGCGGCTGAG | 0.53 | Mid |
| K | GGGCAGCATCGCTGCGAC | 0.33 | Low |
| L | ATTCCGGATGCCGGTTCG | 0.18 | Low |
| M | GATGCTGGGGTTTTGCAT | 0.14 | Low |
| N | AGGGGGGCGGGGAGTGAT | 0.05 | Low |

**Table S4.** β-carotene pathway strain designs and production data.

| UTR tier combinations (n = 27) | | | | | | | | | | |
|---|---|---|---|---|---|---|---|---|---|---|
| Strain ID | UTR Tier (CrtYB-CrtI-CrtE) | UTR ID | | | UTR Strength | | | Combined UTR strength | Beta-carotene (mg/L) | SD |
| | | YB | I | E | YB | I | E | | | |
| S01 | H-H-H | A | B | C | 1.92 | 1.74 | 1.16 | 4.82 | 19.91 | 0.40 |
| S02 | H-H-M | A | B | G | 1.92 | 1.74 | 0.7 | 4.37 | 17.15 | 0.53 |
| S03 | H-H-L | A | B | L | 1.92 | 1.74 | 0.18 | 3.84 | 14.48 | 0.76 |
| S04 | H-M-H | A | G | B | 1.92 | 0.7 | 1.74 | 4.37 | 6.44 | 0.61 |
| S05 | H-M-M | A | G | H | 1.92 | 0.7 | 0.71 | 3.33 | 5.67 | 0.13 |
| S06 | H-M-L | A | G | L | 1.92 | 0.7 | 0.18 | 2.80 | 6.86 | 0.47 |
| S07 | H-L-H | A | L | B | 1.92 | 0.18 | 1.74 | 3.84 | 13.23 | 1.24 |
| S08 | H-L-M | A | L | G | 1.92 | 0.18 | 0.7 | 2.80 | 11.05 | 2.28 |
| S09 | H-L-L | A | L | M | 1.92 | 0.18 | 0.14 | 2.24 | 1.45 | 0.15 |
| S10 | M-H-H | G | A | B | 0.7 | 1.92 | 1.74 | 4.37 | 5.86 | 0.48 |
| S11 | M-H-M | G | A | H | 0.7 | 1.92 | 0.71 | 3.33 | 4.92 | 0.37 |
| S12[a] | M-H-L | G | A | L | 0.7 | 1.92 | 0.18 | 2.80 | 0.15 | 0.00 |
| S13 | M-M-H | G | H | A | 0.7 | 0.71 | 1.92 | 3.33 | 5.10 | 0.44 |
| S14 | M-M-M | G | H | I | 0.7 | 0.71 | 0.64 | 2.05 | 5.04 | 0.54 |
| S15 | M-M-L | G | H | L | 0.7 | 0.71 | 0.18 | 1.59 | 4.89 | 0.49 |
| S16 | M-L-H | G | L | A | 0.7 | 0.18 | 1.92 | 2.80 | 5.77 | 1.40 |
| S17 | M-L-M | G | L | H | 0.7 | 0.18 | 0.71 | 1.59 | 5.06 | 0.50 |
| S18 | M-L-L | G | L | M | 0.7 | 0.18 | 0.14 | 1.02 | 2.79 | 0.18 |
| S19 | L-H-H | L | A | B | 0.18 | 1.92 | 1.74 | 3.84 | 16.58 | 3.02 |
| S20 | L-H-M | L | A | G | 0.18 | 1.92 | 0.7 | 2.80 | 14.61 | 1.25 |
| S21 | L-H-L | L | A | M | 0.18 | 1.92 | 0.14 | 2.24 | 3.82 | 0.25 |
| S22 | L-M-H | L | G | A | 0.18 | 0.7 | 1.92 | 2.80 | 8.38 | 0.78 |
| S23 | L-M-M | L | G | H | 0.18 | 0.7 | 0.71 | 1.59 | 8.09 | 2.71 |
| S24 | L-M-L | L | G | M | 0.18 | 0.7 | 0.14 | 1.02 | 3.57 | 0.42 |
| S25 | L-L-H | L | M | A | 0.18 | 0.14 | 1.92 | 2.24 | 2.57 | 0.08 |
| S26 | L-L-M | L | M | G | 0.18 | 0.14 | 0.7 | 1.02 | 2.50 | 0.04 |
| S27 | L-L-L | L | M | N | 0.18 | 0.14 | 0.05 | 0.37 | 1.00 | 0.15 |

| High-tier UTR permutations (n = 6) | | | | | | |
|---|---|---|---|---|---|---|
| Strain ID | UTR ID (CrtYB-CrtI-CrtE) | UTR Strength | | | Beta-carotene (mg/L) | SD |
| | | CrtYB | CrtI | CrtE | | |
| H1 | A-B-C | 1.92 | 1.74 | 1.16 | 20.60 | 1.37 |
| H2 | A-C-B | 1.92 | 1.16 | 1.74 | 13.70 | 2.48 |
| H3 | B-A-C | 1.74 | 1.92 | 1.16 | 18.62 | 3.11 |
| H4 | B-C-A | 1.74 | 1.16 | 1.92 | 16.06 | 1.27 |
| H5 | C-A-B | 1.16 | 1.92 | 1.74 | 5.60 | 1.19 |
| H6 | C-B-A | 1.16 | 1.74 | 1.92 | 5.67 | 0.72 |

[a]S12 strain showed near-background β-carotene production. [b]yEJ020 (pEJ_S01; UTRb–CrtYB–UTRc–CrtI–UTRa–CrtE) was used as both the S01 strain in the UTR tier combination experiment (Fig. 5B) and the H1 strain in the High-tier permutation experiment (Fig. 5D).

**Table S5.** Primers used in this study.

| Primer name | Purpose | Sequence (5'-3') | Amplicon size |
|---|---|---|---|
| TAF10_F | RT-qPCR reference | CTAAGGATGCCTACGAATATTCCAG | 150 bp |
| TAF10_R | RT-qPCR reference | TCTCATTCTGTTGATGTTGTTGTTG | |
| ZsGreen_F | RT-qPCR target | TGGAGGGAGACTAAGGTGC | 150 bp |
| ZsGreen_R | RT-qPCR target | GCGTGCTCGGTTAAATGC | |
